# A Bio-inspired Latent TGF-β Conjugated Scaffold Improves Neocartilage Development

**DOI:** 10.1101/2025.02.03.636279

**Authors:** Tianbai Wang, Celina C. Maldonado, Bor-Lin Huang, Enkhjargal Budbazar, Andrew Martin, Matthew D. Layne, Joanne E. Murphy-Ullrich, Mark W. Grinstaff, Michael B. Albro

## Abstract

In cartilage tissue engineering, active TGF-β is conventionally supplemented in culture medium at highly supraphysiologic doses to accelerate neocartilage development. While this approach enhances cartilage extracellular matrix (ECM) biosynthesis, it further promotes tissue features detrimental to hyaline cartilage function, including the induction of tissue swelling, hyperplasia, hypertrophy, and ECM heterogeneities. In contrast, during native cartilage development, chondrocytes are surrounded by TGF-β configured in a latent complex (LTGF-β), which undergoes cell-mediated activation, giving rise to moderated, physiologic dosing regimens that enhance ECM biosynthesis while avoiding detrimental features associated with TGF-β excesses. Here, we explore a bio-inspired strategy, consisting of LTGF-β-conjugated scaffolds, providing TGF-β exposure regimens that are moderated and uniformly administered throughout the construct. Specifically, we evaluate the performance of LTGF-β scaffolds to improve neocartilage development with bovine chondrocyte-seeded agarose constructs compared to outcomes from active TGF-β media supplementation (MS) at a physiologic 0.3 ng/mL dose (MS-0.3), supraphysiologic 10 ng/mL dose (MS-10), or TGF-β free. For small-size constructs (∅3×2 mm), LTGF-β scaffolds yield neocartilage that achieves native-matched mechanical properties (800-925 kPa) and sGAG content (6.6%-7.1%), while providing a cell morphology and collagen distribution more reminiscent of hyaline cartilage. LTGF-β scaffolds further afford an optimal chondrogenic phenotype, marked by a 12-to 28-fold reduction of COL-I expression relative to TGF-β-free and a 7-to 17-fold reduction of COL-X expression relative to MS-10. Further, for large-size constructs, which approach the dimensions needed for clinical cartilage repair, LTGF-β scaffolds significantly reduce mechanical and biochemical heterogeneities relative to MS-0.3 and MS-10. Overall, the use of LTGF-β scaffolds improves the composition, structure, material properties, and cell phenotype of neocartilage.

## 1. Introduction

Cartilage tissue engineering is an emerging osteoarthritis treatment strategy for the clinical restoration of degenerated articular cartilage. Here, cell-seeded constructs are cultivated in order to generate neocartilage tissue grafts for surgical implantation in defect sites^1–3^. To achieve this goal, neocartilage needs to recapitulate the composition and structure of the hyaline cartilage extracellular matrix (ECM), comprised of a dense network of sulfated glycosaminoglycans (sGAG) enmeshed in a type-II collagen (COL-II) network, while also maintaining a stable chondrogenic cell population required to provide long-term tissue maintenance^4,5^. To date, tissue engineering protocols are minimally effective, marked by neocartilage that fails to fully recapitulate cartilage composition and that exhibits an unstable chondrogenic cell phenotype, together resulting in a tissue with insufficient clinical survival outcomes^6^. These limitations motivate new efforts to develop the next generation of cell sources, biomaterial scaffolds, and growth factor stimulation protocols to improve clinical outcomes.

Transforming growth factor beta (TGF-β) is a widely used mediator for promoting cartilage regeneration. TGF-β supplemented culture medium can stimulate construct growth during *in vitro* cultivation^7,8^, and when loaded within biomaterial scaffolds (e.g., via affinity domains, liposomes, microspheres) enables sustained stimulation *in vivo* post-implantation^9,10^. Conventionally, TGF-β is administered at supraphysiologic doses, ranging from 10-100 ng/mL for media supplementation protocols^7,8,11^ and 100-10,000 ng/mL for scaffold-delivery protocols^12–14^, representing levels that are one-to-six orders of magnitude higher than TGF-β levels exposed to cells in native cartilage^15^. While supraphysiologic TGF-β doses accelerate ECM biosynthesis, they are also associated with pathogenesis. Administration of supraphysiologic TGF-β doses in synovial joints incites pathology in the form of cellular hyperplasia, hypertrophy, and fibrosis^16–18^. Accordingly, there is a need for caution when administering supraphysiologic TGF-β for cell therapies.

Motivated by these concerns, we recently investigated the effect of moderated, physiologic-range doses of TGF-β on neocartilage development^15^. Contrary to conventional expectations, physiologic doses stimulated sufficient ECM biosynthesis required to produce a functional neocartilage tissue, but did so while mitigating the onset of tissue features detrimental to hyaline cartilage health, including tissue swelling, chondrocyte hyperplasia, and aberrant type-I collagen (COL-I) deposition, thus suggesting that physiologic TGF-β doses may improve neocartilage development. However, despite this promise, physiologic TGF-β doses encounter significant delivery limitations resulting from cell-mediated chemical reactions that rapidly clear TGF-β from within a construct^15,19^. For media supplementation protocols, these reactions limit TGF-β uptake into the construct (Fig.1A), leading to substantial heterogeneities of tissue growth^15,19^. For scaffold delivery platforms, these reactions limit the retention time of loaded TGF-β^20^, affording insufficient growth stimulation. Accordingly, the administration of TGF-β at physiologic doses constitutes a significant challenge.

**Fig 1.**
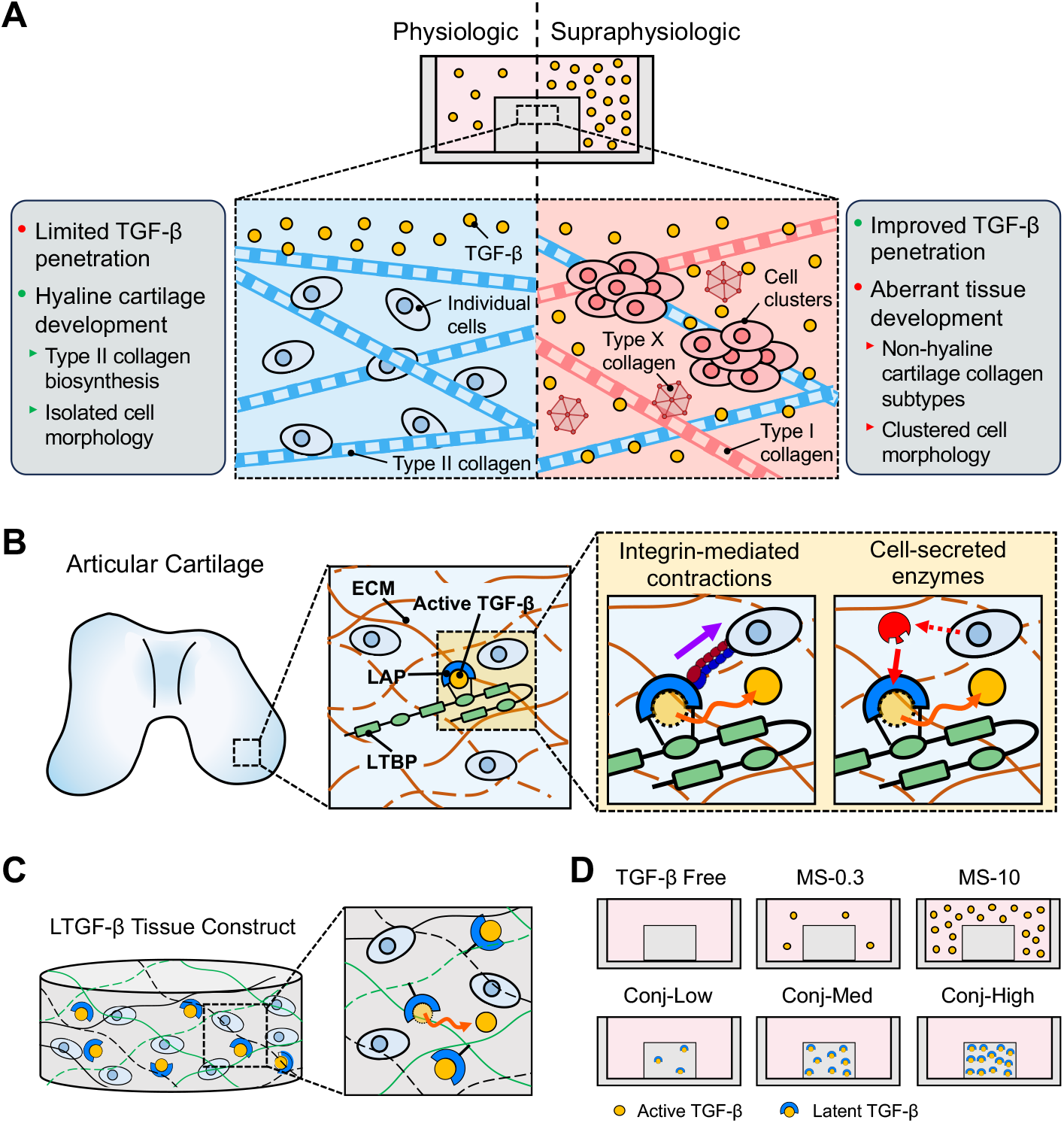
(A) TGF-β delivery paradox in cartilage tissue engineering. Physiologic TGF-β doses (≤0.3 ng/mL) promote hyaline-cartilage-like tissue development but exhibit limited penetration when supplemented in culture medium. Conventionally administered supraphysiologic TGF-β doses (≥10 ng/mL) achieve better penetration into constructs but can induce aberrant tissue features, including non-hyaline cartilage collagen subtypes, cell hyperplasia, and cell hypertrophy. (**B**) **Native TGF-β regulatory processes**. In native cartilage, TGF-β is stored in the ECM in a latent complex (LTGF-β), comprised of sequestration of the active peptide in a latency associated peptide (LAP), which is tethered to ECM via a latent TGF-β binding protein (LTBP). LTGF-β undergoes activation via cell-mediated processes, giving rise to biosynthetic enhancements while avoiding detrimental features associated with TGF-β excesses. (**C**) **Bio-inspired TGF-β delivery platform**. TGF-β is conjugated to a functionalized hydrogel scaffold via coupling of LAP, enabling localized cell-mediated activation. (**D**) **Experimental TGF-β dosing groups**. We evaluate the development of constructs comprised of immature bovine chondrocytes seeded in agarose for up to 56 days in response to variable TGF-β dosing groups: TGF-β-free, physiologic dose media supplementation (0.3 ng/mL; MS-0.3), supraphysiologic dose media supplementation (10 ng/mL; MS-10), and LTGF-β conjugated scaffolds at a low (60 ng/mL; Conj-Low), medium (150 ng/mL; Conj-Med), and high (290 ng/mL; Conj-High) dose.

Interestingly, natural TGF-β regulatory mechanisms have evolved to address this delivery paradox. Here, TGF-β is locally available to cells via an inactive latent complex (latent TGF-β [LTGF-β]), comprised of the active TGF-β peptide immobilized within a propeptide shell (latency associated peptide [LAP]), which in turn is anchored to the ECM via a latent TGF-β binding protein (LTBP) (Fig.1B)^21–23^. Notably, the active TGF-β peptide first needs to be released from the complex, termed activation, to bind to cell receptors and elicit biological responses^22^. Activation occurs via an assortment of triggers, including cell-secreted enzymes^24^, integrin-mediated contraction^25^, thrombospondin-1^26^, and mechanical force transmission^27^. LTGF-β is present at high levels in native cartilage tissue (~300 ng/mL^28^), but activation levels are notably low (less than 1 ng/mL^27,29,30^). This process gives rise to moderated and localized TGF-β exposure regimens, a regulatory dynamic that promotes requisite enhancements of ECM biosynthesis while preventing the induction of tissue pathology that is associated with TGF-β excesses.

Inspired by these native regulatory processes, we describe a hydrogel scaffold incorporated with native levels of the LTGF-β complex to recapitulate natural beneficial regulatory pathways (Fig.1C). This novel platform offers an opportunity to provide moderated and localized TGF-β delivery within a construct via controlled, cell-mediated activation of scaffold-conjugated LTGF-β. We hypothesize that this TGF-β delivery platform will yield: 1) neocartilage with native-matched functional material properties, while mitigating detrimental tissue features associated with supraphysiologic TGF-β dosing, (e.g., tissue swelling, cell, hyperplasia, phenotype dysregulation); and, 2) a more uniform composition for large-size neocartilage constructs as required for clinical chondral repair.

In this study, cartilage regeneration from LTGF-β scaffolds was evaluated in a model system of bovine chondrocytes seeded in a LTGF-β-conjugated agarose hydrogel scaffold. Cartilage regeneration outcomes were compared to those achieved via the administration of active TGF-β in culture media for both a physiologic range dose (0.3 ng/mL) and conventionally-utilized supraphysiologic dose (10 ng/mL) (Fig.1D). In small-size constructs (∅3×2 mm), neocartilage development exhibits only modest spatial gradients of media-supplemented TGF-β, allowing for a faithful comparison of the effect of LTGF-β scaffolds to media-supplemented active TGF-β in the absence of spatial distribution artifacts^15^ as assessed by mechanical properties, ECM composition, cell morphology, and gene expression profiles. Subsequently, we evaluated the ability of LTGF-β scaffolds to mitigate developmental heterogeneities on larger-size constructs (∅5×2 mm), which approach the dimensions of neocartilage grafts required for clinical cartilage repair.

## 2. Materials and methods

### 2.1. TGF-β dosing groups

Bovine chondrocyte-seeded agarose tissue constructs were prepared at a small-size (∅3×2 mm) and large-size (∅5×2 mm) configuration. Constructs were cultivated with LTGF-β1-conjugated agarose scaffolds over a range of conjugation levels: low (Conj-Low), medium (Conj-Med) and high (Conj-High) levels (preparation details described below). Development outcomes were compared to constructs cultivated with media supplementation (MS) of active TGF-β3 in culture medium for the initial 14 days of culture at a physiologic (0.3 ng/mL; MS-0.3) or supraphysiologic (10 ng/mL; MS-10) dose, or maintained entirely without TGF-β (TGF-β-free) (Fig.1D). While different TGF-β isoforms were used for active TGF-β media supplementation and LTGF-β scaffold conjugation, we did not observe isoform-dependent differences in construct development for bovine chondrocytes (Fig.S1). For all experiments, constructs were cultivated in chondrogenic media consisting of high-glucose Dulbecco’s Modified Eagle Medium (hgDMEM), supplemented with 1% ITS+ premix (Corning), 1 mM sodium pyruvate, 50 μg/mL L-proline, 1% PS/AM antibiotic-antimycotic, 100 nM dexamethasone, and 50 μg/mL ascorbate-2-phosphate at 37°C at a 40:1 media to tissue volume ratio with media refreshed three times per week. Small-size constructs were evaluated for: 1) mechanical properties after 28 and 56 days of culture, 2) cell morphology after 42 days, and biochemical content, histology, and immunostaining after 56 days of culture, and 3) RT-PCR after 14, 28, and 45 days of culture. Large-size constructs were evaluated for: 1) bulk mechanical properties after 28 and 56 days of culture, 2) spatial mechanical analysis after 56 days of culture, 3) bulk biochemical content and histology after 56 days of culture.

### 2.2. Methacrylate agarose synthesis and LTGF-β conjugation

Five grams of type VII agarose (Agr; Sigma) was dissolved in 150 mL DMSO at 120°C. After cooling to 60°C, 8 mg of 4-dimethylaminopyridine, 16 mg of hydroquinone, and 500 μL of methacrylic anhydride were reacted for 1 hour under continuous stirring^31^. Methacrylate agarose (MeAgr) was acetone-precipitated, minced, washed in acetone, suspended in deionized water, and lyophilized. From preliminary assessments, MeAgr did not have a detrimental effect on chondrocyte ECM biosynthesis (Fig.S2). Human recombinant LTGF-β1 (small latent complex [SLC] form, R&D Systems) was conjugated to agarose via incubation in 4% w/v MeAgr at 37°C overnight at the following levels prior to cell encapsulation: 2.5 μg/mL (Conj-Low), 6 μg/mL (Conj-Med), or 12 μg/mL (Conj-High).

### 2.3. Tissue construct fabrication

Articular cartilage was harvested from eight bovine carpometacarpal joints (3-6 months old). Cartilage was digested for 16 hours at 37°C with 1000 U/mL type IV collagenase (Worthington) supplemented in hgDMEM with 5% fetal bovine serum (FBS)^32^. Isolated chondrocytes were suspended in hgDMEM at 60×10^6^ cells/mL and mixed with LTGF-β1-conjugated 4% w/v MeAgr and unmodified 4% w/v Agr in a 2:1:1 volume ratio to achieve a final mixture of 30×10^6^ cells/mL in a 2% w/v MeAgr/Agr blend (Fig.2A).

**Fig 2.**
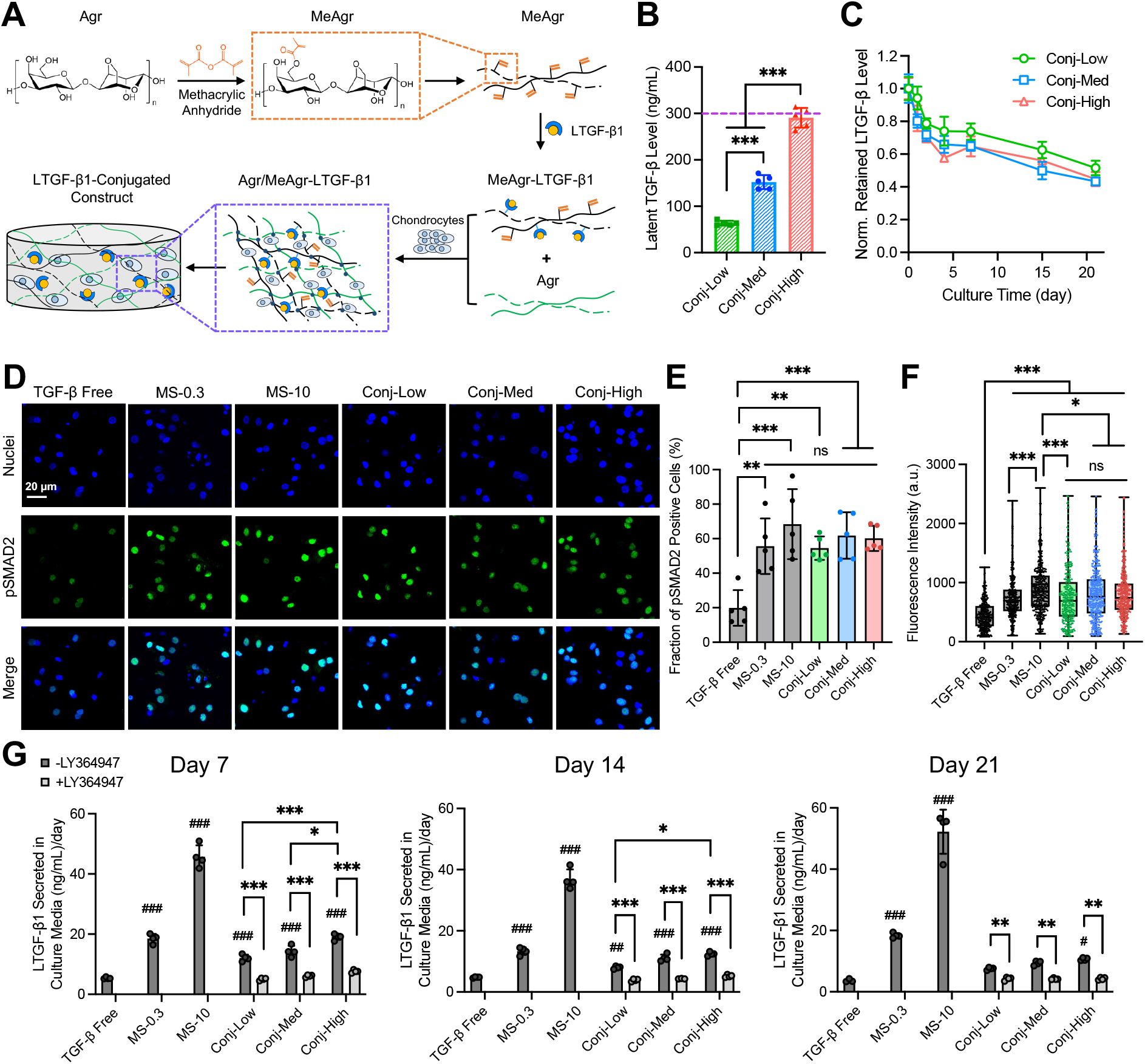
LTGF-β scaffold conjugation yields and bioactivity. (**A**) Agarose (Agr) is reacted with methacrylic anhydride for methacrylate agarose (MeAgr) synthesis. MeAgr is conjugated with LTGF-β1, mixed with Agr in a 1:1 ratio, and embedded with bovine chondrocytes to fabricate tissue constructs. (**B**) Initial levels of conjugated LTGF-β1 in constructs for Conj-Low, Conj-Med, and Conj-High doses. n=5. (**C**) Retention profiles of conjugated LTGF-β1 in constructs over 21 days. n=5. (**D**) Representative immunostaining images of pSMAD2 and nuclei for TGF-β-free, MS-0.3, MS-10, and Conj-Low, Conj-Med, and Conj-High doses at day 7. n=5. (**E**) Fraction of pSMAD2 positive cells and (**F**) mean pSMAD2 fluorescence intensity in cell nuclear regions. (**G**) LTGF-β1 secretion levels in conditioned media at day 7, 14, and 21, in the absence or presence of a TGF-β receptor kinase inhibitor, LY364947. n=4. Error bars represent mean ± s.d. *p<0.05, **p<0.005, ***p<0.001; ^##^p<0.005, ^###^p<0.001: significant difference to the TGF-β Free levels; ns: no significance.

### 2.4. LTGF-β conjugation levels and retention profiles

∅3×2 mm LTGF-β constructs for all conjugated doses were collected at days 0, 1, 2, 4, 7, 14, and 21 to evaluate the LTGF-β1 retention profiles. Here, constructs were initially treated overnight with a Tris/CHAPS buffer solution (50 mM Tris/HCl, 5% CHAPS), which exclusively extracts unconjugated LTGF-β from the construct^33^. Subsequently, constructs were treated overnight with 4M guanidine hydrochloride at 4°C, which activated and extracts scaffold-conjugated LTGF-β. After each treatment step, extracts were diluted in 1 mg/mL BSA solution and analyzed via a TGF-β1 ELISA (Duoset, R&D Systems), as described previously^19,33^.

### 2.5. Autoinduction-based TGF-β activity assessment

To evaluate the activity level of conjugated LTGF-β, our previously developed autoinduction-based quantification platform was used where TGF-β cell exposure levels are inferred from the analysis of newly synthesized endogenous LTGF-β^30^. Conj-Low, Conj-Med, and Conj-High constructs were cultured for 21 days with media replenished three times per week. Conditioned media were collected at day 7, 14, and 21. The level of LTGF-β1 in conditioned media was measured using an isoform-specific TGF-β1 ELISA (Duoset, R&D Systems). To confirm that LTGF-β1 secretion enhancements resulted from activation of scaffold-conjugated LTGF-β, additional constructs were cultured in the presence of LY364947, a TGF-β receptor kinase inhibitor. TGF-β-free, MS-0.3, and MS-10 constructs were cultured in parallel for reference.

### 2.6. Mechanical property assessments

For bulk property measures, constructs were subjected to a 10% axial strain in unconfined compression. The Young’s modulus (E_Y_) was calculated based on the measured stress at equilibrium. The mechanical heterogeneities of ∅5×2 mm constructs were characterized via digital image correlation (DIC) analysis^15,34^. Here, constructs were diametrically halved and cell nuclei were stained with 1 mg/mL Hoechst 33342 (Thermo Scientific). The exposed cross-section was imaged via confocal microscopy (Olympus FV3000, 10×) before and after application of a 10% platen-to-platen compressive strain. The 2D strain distribution was assessed using Ncorr (Georgia Institute of Technology) and binned into six sections of uniform thickness through the construct depth. For each tissue, the mean strain of each section was normalized to the mean strain in the media-exposed topmost tissue section. A linear regression was applied to the curve of normalized strain versus section number, and the resulting slope was designated as strain gradient.

### 2.7. Biochemical content analysis

Constructs were digested overnight in 0.5 mg/mL Proteinase K at 56°C (Fisher BioReagents). Tissue digests were assayed for DNA content via PicoGreen dsDNA assay (Invitrogen) and sGAG via dimethylmethylene blue assay^35^. Tissue digests were further hydrolyzed and assayed for collagen content via orthohydroxyproline assay^36^. sGAG and collagen contents were presented as mass per construct, mass per construct wet weight (% ww), and mass per DNA content.

### 2.8. Cell morphology analysis

Constructs were diametrically halved, stained with Live/Dead kit (Invitrogen), and the exposed cross section was imaged via confocal microscopy (Olympus FV3000, 20×). Cell morphology was evaluated by identifying isolated and clustered cells using a custom algorithm as detailed in our prior work^15^. Briefly, images were binarized and a morphology factor was defined according to the area, circularity, and fluorescence variation of the identified cellular regions to determine isolated and clustered cell morphology. For each construct, results were presented as a cluster area fraction (CAF), defined as the area of clustered cellular regions normalized to the total cellular area.

### 2.9. Histology and immunofluorescence staining

Constructs were fixed in 3.7% buffered formaldehyde at 4°C overnight. For pSMAD2 staining, ∅2×2 mm day 7 constructs were diametrically halved, permeabilized with 1% Triton X-100 and 0.1 g/mL sucrose, blocked with 2% BSA, and incubated with rabbit antibody against phosphorylated-SMAD2 (pSMAD2; 1:100, MBS9211547, MyBioSource) followed by anti-rabbit Alexa 488 (1:200, Invitrogen) and mounting (ProLong™ Gold Antifade, Invitrogen). For collagen subtype staining, 80 μm sections from ∅3×2 mm day 56 constructs were prepared following previous work^37^ and treated with 0.1 mg/mL pronase, blocked, and incubated with mouse antibody against COL-II (1:100, CIIC1, Developmental Studies Hybridoma Bank) and rabbit antibody against type VI collagen (COL-VI; 1:500, ab6588, Abcam), followed by anti-mouse Alexa 594 and anti-rabbit Alexa 488 (1:200, Invitrogen) and mounting. For sGAG staining, 7 μm sections from paraffin blocks were stained with Safranin O/Fast green. Immunostaining images were acquired using confocal microscopy (Olympus FV3000, 40× for pSMAD2, 20× for collagen subtypes) and histology images were acquired using a slide scanner (Olympus VS120, 10×).

### 2.10. Gene expression analysis

Constructs RNA was extracted via RNeasy Plant Mini Kit (Qiagen)^38^ and reverse transcribed into cDNA using High-Capacity cDNA Reverse Transcription Kit (Applied Biosystems). Gene expression was measured via real-time quantitative PCR (RT-PCR) using TaqMan Fast Advanced Master Mix (Applied Biosystems) and TaqMan assay probes for beta-actin (ACTB, Bt03279174_g1), COL-II, (COL2A1, Bt03251861_m1), aggrecan (ACAN, Bt03212186_m1), type-X collagen (COL-X; COL10A1, Bt03215582_m1), COL-I (COL1A1, Bt01463861_g1), SOX9 (Bt07108872_m1), and RUNX3 (Bt04315916_m1). Relative gene expression levels were analyzed by 2^-ΔΔCt^ method, internally normalized to beta-actin, and compared to day 14 TGF-β free group.

### 2.11. Statistical analysis

All experiments were conducted using at least three replicates per condition. Two-way ANOVAs (α = 0.05) were performed to determine the effect of: 1) LTGF-β1 conjugation dose and LY364947 supplementation on LTGF-β levels in conditioned media (Fig.2G), 2) TGF-β dosing group and culture duration on Young’s modulus (Fig.4A and Fig.7A), and 3) TGF-β dosing group and culture duration on gene expression levels (Fig.5). One-way ANOVAs (α = 0.05) were performed to determine the effect of 1) LTGF-β1 administered dose on LTGF-β1 conjugation levels (Fig.2B), 2) LTGF-β1 conjugation dose on pSMAD2 level (Fig.2E-F), 3) TGF-β dosing group on biochemical contents (Fig.3 and Fig.7B-J), 4) TGF-β dosing group on cell cluster morphology (Fig.6B), and 5) TGF-β dosing group on tissue strain gradient (Fig.7N). Tukey’s HSD post-hoc tests were run to examine differences between experimental groups.

**Fig 3.**
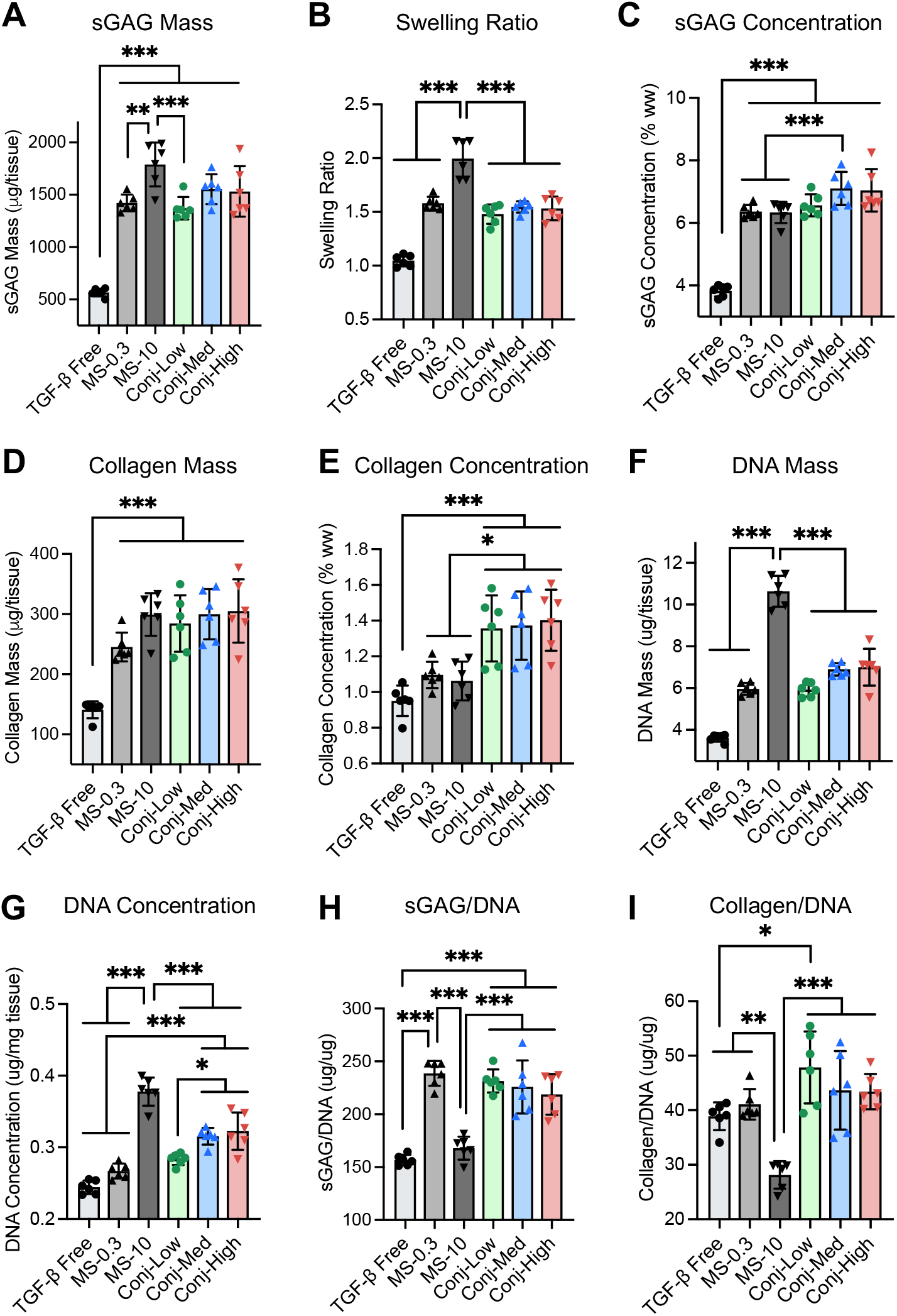
Biochemical content of small-size constructs. (**A**) Mass of sGAG, (**B**) swelling ratio, (**C**) concentration of sGAG, (**D**) mass of collagen, (**E**) concentration of collagen, (**F**) mass of DNA, (**G**) concentration of DNA, (**H**) sGAG-to-DNA mass ratio, (**I**) and collagen-to-DNA mass ratio of ∅3×2 mm constructs at day 56 for TGF-β-free, MS-0.3, MS-10, Conj-Low, Conj-Med, and Conj-High TGF-β dosing groups. n=6. Error bars represent mean ± s.d. *p<0.05, **p<0.005, ***p<0.001.

**Fig 4.**
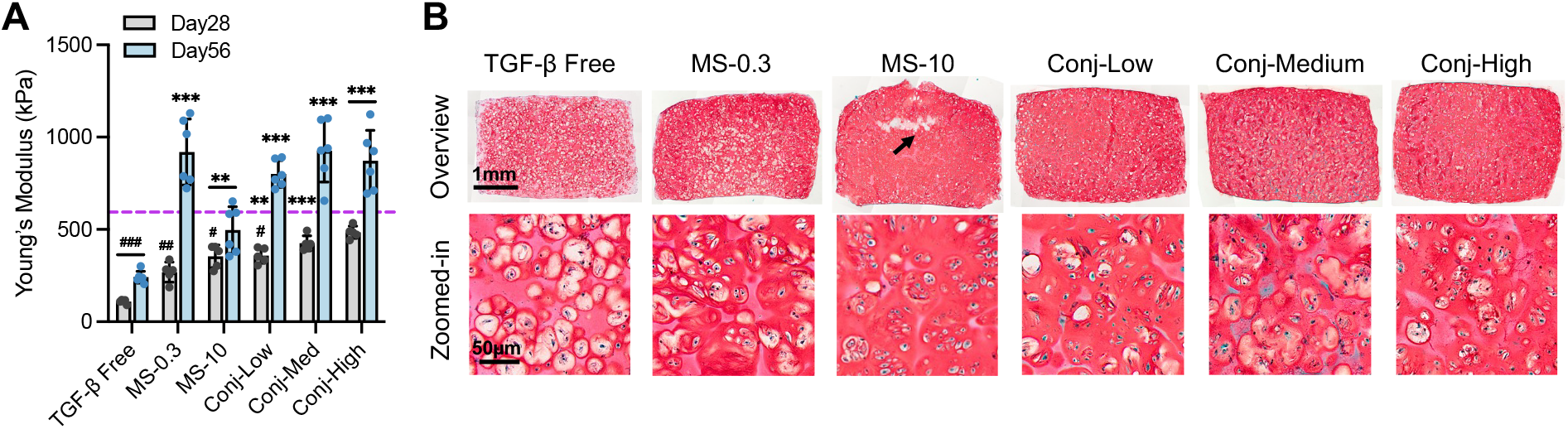
Mechanical properties and histology of small-size constructs. **(A)** Young’s modulus at days 28 and 56 (n=5), and **(B)** Safranin O staining at day 56 (n=3) from ∅3×2 mm constructs for TGF-β-free, MS-0.3, MS-10, Conj-Low, Conj-Med, and Conj-High TGF-β dosing groups. Dashed line represents native cartilage Young’s modulus levels. Arrow depicts construct fissure. Error bars represent mean ± s.d. *p<0.05, **p<0.005, ***p<0.001: significant difference to TGF-β-free levels for the corresponding time points; ^#^p<0.05, ^##^p<0.005, ^###^p<0.001: significantly lower than native levels.

**Fig 5.**
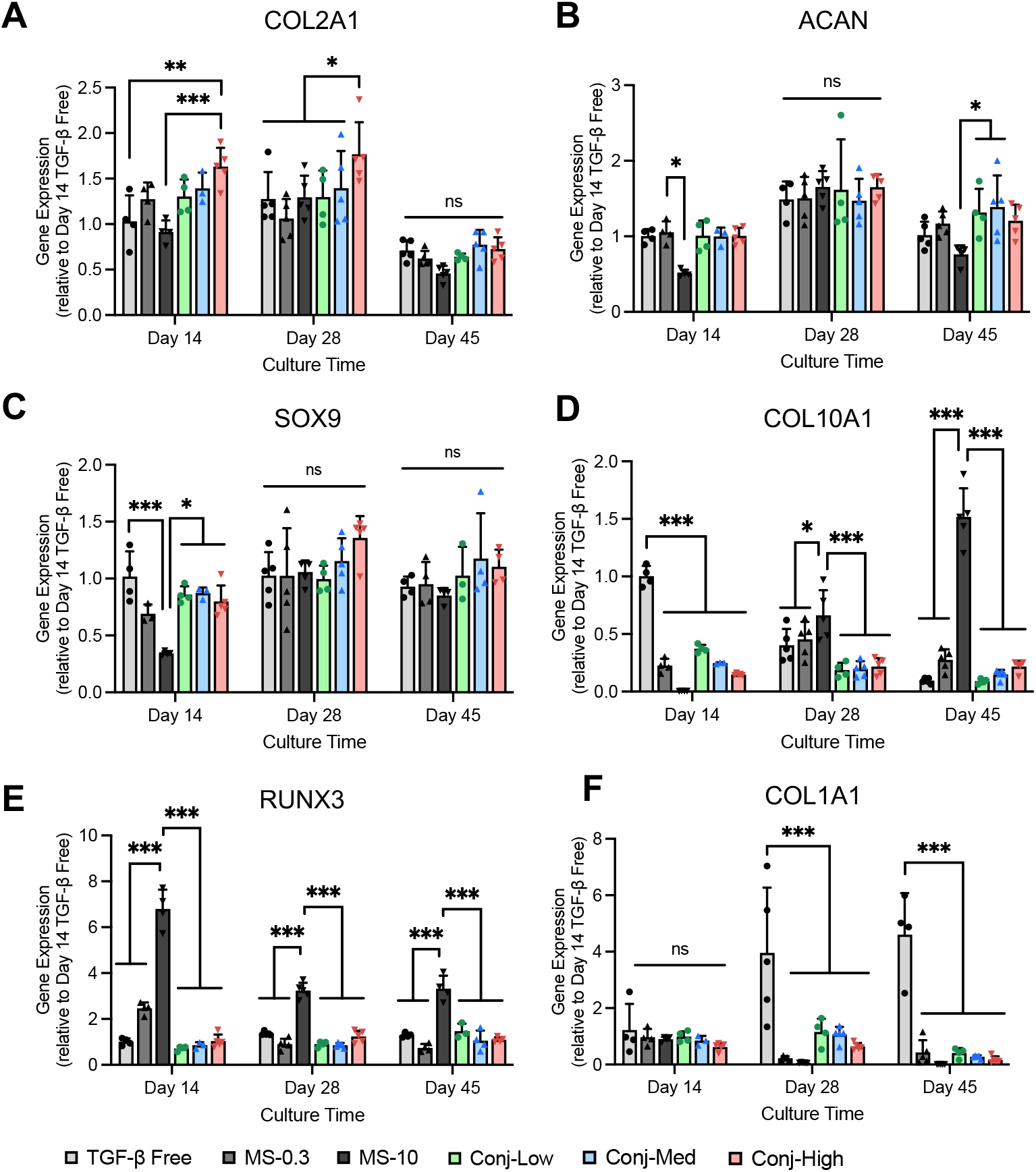
Gene expression for small-size constructs. Gene expression levels of (**A**) COL-II, (**B**) aggrecan, (**C**) SOX9, (**D**) COL-X, (**E**) RUNX3, and (**F**) COL-I in ∅3×2 mm constructs at days 14, 28, and 45 for TGF-β-free, MS-0.3, MS-10, Conj-Low, Conj-Med, and Conj-High TGF-β dosing groups. n=5. Error bars represent mean + s.d. ns: no significance. *p<0.05, **p<0.005, ***p<0.001.

**Fig 6.**
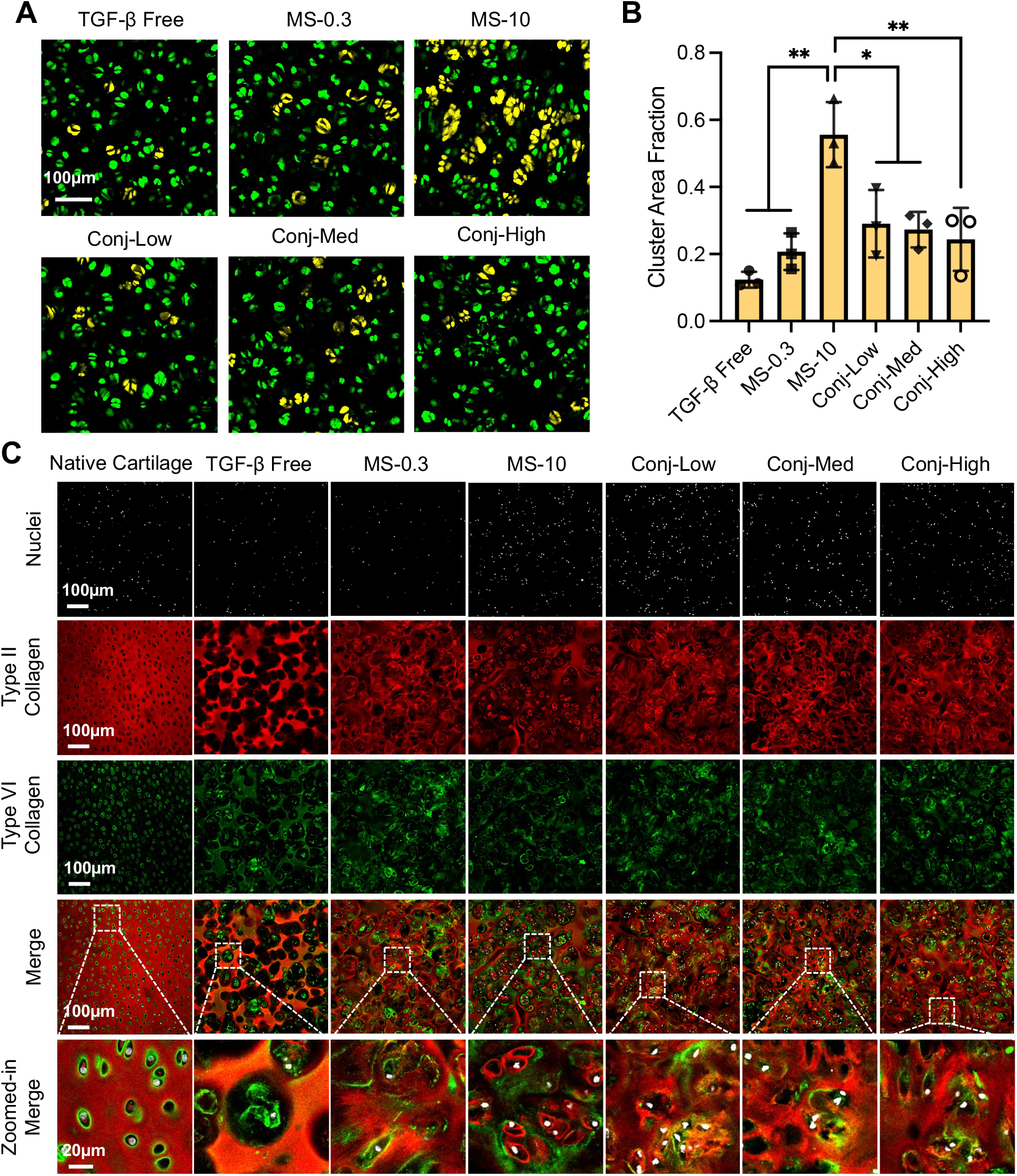
Chondrocyte morphology and ECM distribution in small-size constructs. **(A)** Calcein AM staining for cell morphology in ∅3×2 mm constructs at day 42 for TGF-β-free, MS-0.3, MS-10, Conj-Low, Conj-Med, and Conj-High TGF-β dosing groups. Green: individual cells; Yellow: clustered cells. n=3. (**B**) Cell cluster area fraction (CAF) of constructs quantified from the cell morphology images. Error bars represent mean ± s.d. *p<0.05, **p<0.005, ***p<0.001. (**C**) Immunostaining of COL-II and COL-VI for TGF-β-free, MS-0.3, MS-10, Conj-Low, Conj-Med, and Conj-High TGF-β dosing groups at day 56. n=3.

**Fig 7.**
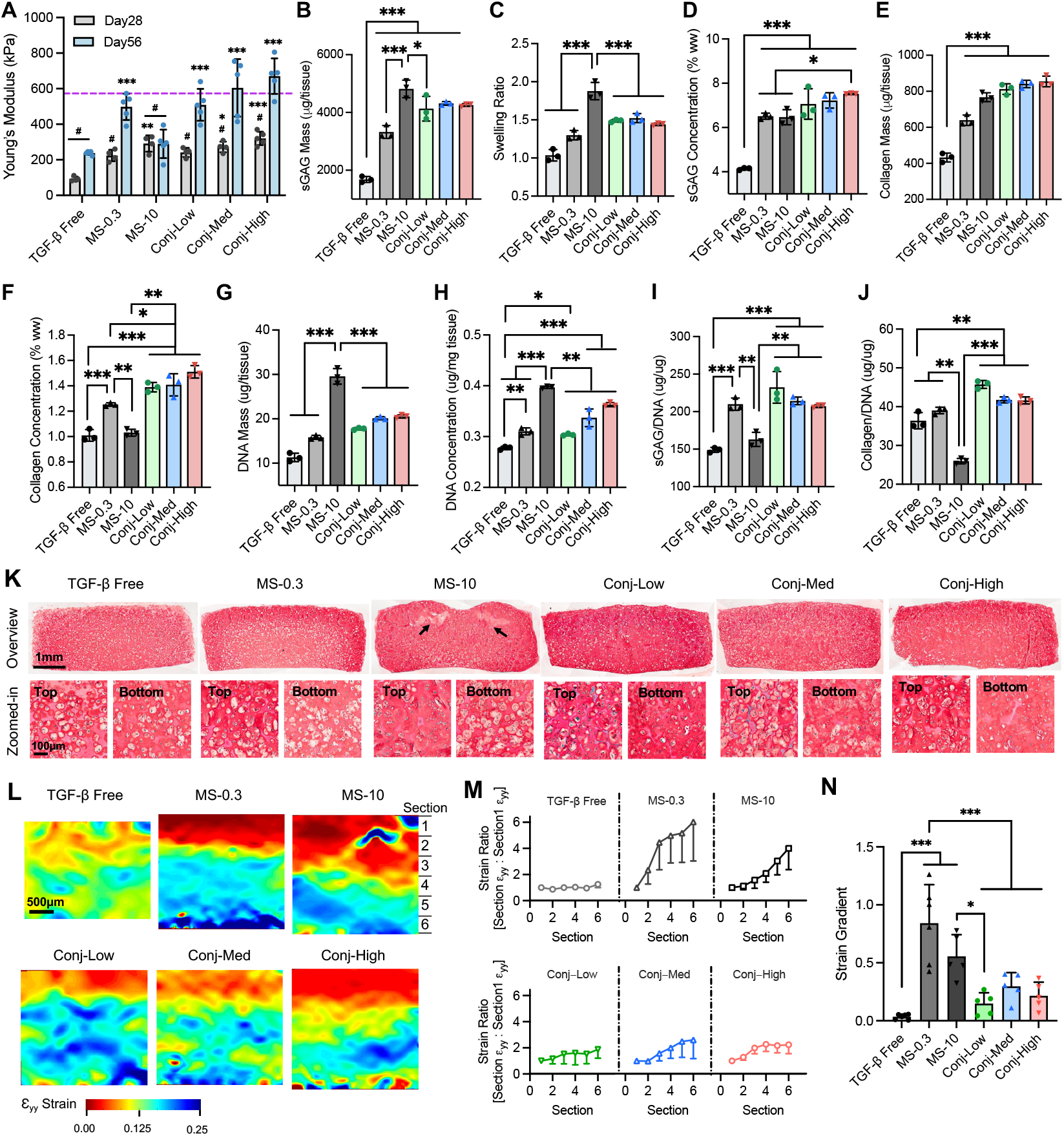
Bulk and spatial biochemical content and mechanical properties in large-size constructs. (**A**) Young’s modulus at days 28 and 56 from ∅5×2 mm constructs for TGF-β-free, MS-0.3, MS-10, Conj-Low, Conj-Med, and Conj-High TGF-β dosing groups. Dashed line represents native cartilage Young’s modulus levels. n=5. Error bars represent mean ± s.d. *p<0.05, **p<0.005, ***p<0.001: significant difference to TGF-β-free levels for the corresponding time points; ^#^p<0.05: significantly lower than native levels. (**B**) Mass of sGAG, (**C**) swelling ratio, (**D**) concentration of sGAG, (**E**) mass of collagen, (**F**) concentration of collagen, (**G**) mass of DNA, (**H**) concentration of DNA, (**I**) sGAG-to-DNA mass ratio, and (**J**) collagen-to-DNA mass ratio for day 56 constructs. n=3. Error bars represent mean ± s.d. (**K**) Full construct section (top row) and high magnification Safranin O staining images of the top and bottom regions. n=3. Arrows depict construct fissure. (**L**) Representative optically-measured compressive strain (ε_yy_) distribution in the central regions of in ∅5×2 mm day 56 constructs in response to 10% platen-to-platen compression. n=5. (**M**) Normalized average strain of each depth-dependent tissue section. Error bars represent mean − s.d. (**N**) Depth-dependent strain gradient (slope of strain ratio) of tissue constructs. Error bars represent mean + s.d. *p<0.05, **p<0.005, ***p<0.001.

## 3. Results

### 3.1. Scaffolds exhibit native LTGF-β content and physiologic bioactivity

Scaffold-conjugated LTGF-β levels increased with initial LTGF-β loading dose, yielding concentrations of 64±5, 152±15, and 290±21 ng/mL for Conj-Low, Conj-Med, and Conj-High groups, respectively (Fig.2B). Conjugated LTGF-β levels decreased gradually over time in culture, but a large fraction (43-52% of the initial dose) remained in the construct after 3 weeks (Fig.2C). LTGF-β conjugation had no effect on cell viability over 42 days of culture (Fig.S3). To evaluate bioactivity of scaffold-conjugated LTGF-β, we assessed TGF-β signaling downstream events including SMAD2 phosphorylation^39^ and TGF-β autoinduction response^30^. The number of pSMAD2 positive chondrocytes and pSMAD2 intensity (Fig.2D-F) in all LTGF-β conjugation groups were significantly higher than TGF-β-free constructs (p<0.005) and similar to MS-0.3 levels, suggesting that chondrocytes were able to activate scaffold-conjugated LTGF-β *in situ*. Further, we observed an autoinduction response in LTGF-β conjugated constructs, whereby the secretion of endogenous LTGF-β1 into conditioned medium was elevated relative to TGF-β-free constructs (p<0.05; Fig.2G), thus further supporting that chondrocytes were able to activate exogenous scaffold-conjugated LTGF-β *in situ*. LTGF-β1 secretion levels were significantly decreased in the presence of LY364947 (p<0.005), confirming that this secretion resulted from enhanced synthesis of endogenous LTGF-β1, rather than desorption of exogenous LTGF-β1 from the scaffold.

### 3.2. LTGF-β scaffolds yield constructs with improved ECM composition

Physiologic TGF-β dosing groups (MS-0.3 and all LTGF-β doses) significantly enhanced the mass of sGAG deposited in small-size constructs, relative to TGF-β-free (p<0.001; Fig.3A). MS-10 enhanced sGAG mass deposition further (p<0.005), indicating even greater sGAG biosynthesis for this supraphysiologic dose. However, MS-10 additionally introduced a pronounced tissue swelling response, marked by an ~2-fold increase in construct weight above day 0 levels (Fig.3B). This swelling was significantly mitigated by physiologic TGF-β dosing groups (MS-0.3 and LTGF-β doses) (p<0.001). Due to this differential swelling behavior, the resulting concentration of sGAG per construct weight is similar between all TGF-β groups (Fig.3C) and all TGF-β groups exhibited native-matched sGAG content (>6% w/v). Similarly, while MS-10 enhanced collagen mass (p<0.001; Fig.3D), the concentration of collagen per construct wet weight was not significantly higher than TGF-β-free (p=0.76; Fig.3E), as a result of dilution of deposited collagen from tissue swelling. In contrast, all LTGF-β doses yielded enhanced collagen content, relative to TGF-β-free and MS-10 (p<0.001 and p<0.05, respectively). MS-10 also induced pronounced cell proliferation, marked by a 3-fold enhancement of DNA mass (p<0.001; Fig.3F) and a 1.5-fold enhancement of DNA concentration (p<0.001; Fig.3G), relative to TGF-β-free. This proliferation was mitigated by physiologic TGF-β dosing groups (MS-0.3 and LTGF-β doses), marked by a significantly lower DNA content relative to MS-10 (p<0.001). Due to this moderate cell proliferation, LTGF-β and MS-0.3 groups exhibited enhanced ECM content on a per cell basis, relative to MS-10, as marked by elevated sGAG (p<0.001; Fig.3H) and collagen per DNA (p<0.001; Fig.3I).

### 3.3. LTGF-β scaffolds yield constructs with native-matched mechanical properties

For small-size constructs, all TGF-β groups induced a greater than 2.5-fold increase in Young’s modulus (E_Y_) at day 28, relative to TGF-β-free (Fig.4A; p<0.05). Notably, all TGF-β groups achieved further E_Y_ enhancements at day 56 except for MS-10. This cessation of tissue growth for MS-10 appears to result from the formation of a fissure in the center of the construct (Fig.4B, top), which leads to compromised compressive tissue properties. Tissue cracking could result from rapid acceleration of sGAG deposition in the absence of a mature collagen network to provide tensile restraint. In contrast, physiologic TGF-β dosing groups (MS-0.3 and LTGF-β doses) significantly enhanced E_Y_ above TGF-β-free and MS-10 levels at day 56 (p<0.001), yielding constructs with native-cartilage-matched mechanical properties.

### 3.4. LTGF-β scaffolds yield constructs with a more chondrogenic cell phenotype

All TGF-β groups exhibited similar levels of COL-II and ACAN expression in small-size constructs (Fig.5A-B). MS-10 induced a significant reduction in expression of the chondrogenic transcription factor, SOX9, at day 14 (p<0.001; Fig.5C), but its expression levels recovered for subsequent time points after removal of this supraphysiologic TGF-β dose from culture. MS-10 also significantly increased expression of COL-X (Fig.5D), a hypertrophic biomarker, at days 28 and 45 (p<0.005), and RUNX3 (Fig.5E), a hypertrophic transcription factor, at all time points, relative to TGF-β-free (p<0.001). In contrast, physiologic TGF-β dosing groups (MS-0.3 and LTGF-β doses) maintained COL-X and RUNX3 at TGF-β-free levels. Further, COL-I expression was elevated at days 28 and 45 for TGF-β-free constructs (Fig.5F). All TGF-β groups significantly reduced Col-I expression (p<0.001), indicating that physiologic TGF-β dosing is sufficient to suppress expression of this non-hyaline-cartilage-associated collagen subtype.

### 3.5. LTGF-β scaffolds yield constructs with improved cell morphology and ECM distribution

TGF-β-free groups maintained chondrocytes in an isolated, spherical morphology, as exhibited by a low CAF value (0.12±0.02; Fig.6A-B), consistent with the characteristic cell morphology of healthy hyaline cartilage^15^. MS-10 yielded constructs with cells in a highly clustered morphology, as exhibited by a significant increase in CAF (0.56±0.10), consistent with the induction of high cell proliferation rates (Fig.3F-G). Physiologic TGF-β dosing (MS-0.3 and LTGF-β doses) significantly mitigated cell clustering (p<0.05), yielding constructs with CAF values less than 0.3.

LTGF-β scaffolds further yielded a more hyaline cartilage-like ECM distribution. The native hyaline cartilage structure is characterized by a thin ring of COL-VI in the pericellular matrix (PCM), and an abundance of COL-II in surrounding territorial and interterritorial regions (Fig.6C). For TGF-β-free, the territorial region was devoid of COL-II. For MS-10, COL-II was localized to the PCM but devoid in the interterritorial matrix. In contrast, for physiologic TGF-β dosing (MS-0.3 and LTGF-β doses), COL-II was present in both territorial and interterritorial regions, closer to the distribution characteristic of native hyaline cartilage.

### 3.6. LTGF-β scaffolds yield large-size constructs with reduced ECM heterogeneities

Large-size constructs exhibited trends of bulk mechanical properties and biochemical contents similar to small-size constructs (Fig.7A-J). Notably, LTGF-β constructs reached native-matched Young’s modulus (Fig.7A) and sGAG concentration (Fig.7D) levels and significantly mitigated swelling relative to MS-10 (p<0.001; Fig.7C). Further, LTGF-β constructs significantly mitigated cell proliferation relative to MS-10 (p<0.005; Fig.7G-H), yielding enhanced ECM content on a per cell basis (p<0.005; Fig.7I-J). For spatial properties, TGF-β-free constructs exhibited a relatively homogeneous distribution of sGAG deposition and mechanical properties, as exhibited by near uniform Safranin O staining (Fig.7K) and a uniform depth-dependent strain distribution in response to axial compression (strain gradient =0.03±0.02; Fig.7L-N). Media supplemented active TGF-β (MS-0.3 and MS-10) induced substantial heterogeneities in tissue composition and mechanical properties. MS-0.3 constructs exhibited heterogeneous sGAG distribution, with Safranin O intensity decreasing with distance from the media-exposed construct surface. MS-10 constructs exhibited macroscale fissuring beneath the construct surface, consistent with a spatially varied Donnan-osmotic swelling pressure that results from heterogeneities of sGAG. MS-0.3 and MS-10 further exhibited pronounced heterogeneities in mechanical properties, as exhibited by a non-uniform depth-dependent strain distribution in response to axial compression (strain gradient =0.84±0.33 and 0.56±0.19, respectively) In contrast, LTGF-β scaffolds mitigated these biochemical and mechanical heterogeneities. All LTGF-β doses exhibited a near uniform sGAG distribution without tissue fissures. Further, LTGF-β constructs exhibited more uniform strain distributions, as evidenced by strain gradient levels significantly lower than MS-0.3 (strain gradient=0.15±0.09, 0.30±0.12, 0.22±0.12, for Conj-Low, Conj-Med, Conj-High, respectively; p<0.001).

## 4. Discussion

A long-standing challenge in cartilage tissue engineering is the generation of neocartilage that recapitulates the composition, structure, and material properties required for cartilage function, while maintaining a stable chondrogenic phenotype needed to maintain long-term tissue health. The use of supraphysiologic doses of active TGF-β is a conventional strategy to accelerate ECM biosynthesis and promote chondrogenesis. However, as illustrated previously^15^ and in this study, this strategy encounters significant drawbacks, which include the induction of tissue swelling, cell hyperplasia, aberrant cell phenotypes, and ECM heterogeneity. To address these limitations, we explored an alternative bio-inspired strategy, consisting of TGF-β delivery via scaffold conjugation in its natural latent complex, providing moderate and localized TGF-β exposure regimens throughout the construct, akin to what occurs in native cartilage. Here we show that the LTGF-β scaffold can improve the composition, structure, material properties, and cell phenotype of neocartilage.

LTGF-β has been used to promote cartilage regeneration in prior studies^40,41^. However, this is the first study to evaluate the effect of LTGF-β on neocartilage development when administered at levels present in native cartilage (~300 ng/mL^28^). Chemical conjugation of LTGF-β, in its small latent complex form, to methacrylate functionalized agarose via Michael addition of lysine residues on the LAP shell region afford the LTGF-β scaffold (Fig.2A). Conjugated LTGF-β undergoes a slow release from the scaffold likely due to hydrolysis of the ester bond linking LAP to agarose. However, ~50% of conjugated LTGF-β remains after 21 days of culture, thus providing a stable reservoir of LTGF-β to cells. Notably, this molecular configuration is distinct from that presents in native cartilage, whereby the cell-secreted LTGF-β anchors the ECM through an intermediate LTGF-β binding protein (LTBP), which acts to modulate matrix incorporation and subsequent activation processes^23,42^ (Fig.1B). Despite these structural differences, conjugated LTGF-β remains accessible to chondrocytes for activation, as evidenced by positive pSMAD2 signaling in LTGF-β constructs (Fig.2D-F). Activation likely results from cell-mediated activation mechanisms that occur in native tissue, such as integrin-mediated cell traction or cell-secreted enzymes^21,24,25^. While quantitative measures of LTGF-β activation levels are difficult to obtain, autoinduction assessments suggest that LTGF-β scaffolds provide cells with TGF-β doses <0.3 ng/mL (Fig.2G), consistent with the range of physiologic levels present in native cartilage^27,29,30^. Interestingly, all LTGF-β dosing groups induce similar construct development outcomes, suggesting that a range of physiologic doses are sufficient to induce neocartilage development. In addition, enhancements of ECM biosynthesis are not present when LTGF-β is administered in non-functionalized agarose scaffolds (Fig.S4), suggesting that conjugation to the scaffold is required for LTGF-β activation to occur.

The generation of functional neocartilage requires the deposition of native levels of ECM constituents to yield tissue properties sufficient to support mechanical loads. Supraphysiologic doses of active TGF-β have traditionally been adopted to accelerate sGAG deposition and material property development^7,8,28^. Consistent with this expectation, constructs exposed to supraphysiologic TGF-β exhibit significantly enhanced sGAG deposition (Fig.3A&C), yielding a near-native compressive Young’s modulus after only 1 month of culture (Fig.4A). However, this accelerated tissue development is accompanied by a detrimental tissue swelling response, whereby constructs swell by ~200% of their initial volume (Fig.3B). This behavior arises from a sGAG-induced Donnan-osmotic swelling pressure, which is unrestrained due to the slow-to-develop collagen tensile network^43,44^. Tissue swelling initially dilutes the construct collagen content reducing tensile properties (Fig.3D-E) and eventually leads to a large-size fissure in the construct center (Fig.4B), which compromises the tissue’s capacity to support mechanical loads^45^. In contrast, physiologic TGF-β dosing via LTGF-β scaffolds promotes sGAG biosynthesis at a more measured rate, yielding the deposition of a native-levels of sGAG while mitigating tissue swelling. This enables the generation of neocartilage that rapidly recapitulates functional compressive properties while avoiding tissue fissuring.

Neocartilage functionality further depends on the spatial organization and structure of the tissue, as required for transmitting mechanical loads between cells and ECM, which regulates mechanochemical signaling cues^46^. The arrangement of cells in dense clusters is a striking feature of neocartilage stimulated with supraphysiologic TGF-β (Fig.6A-B), given their morphological similarity to pathologic clustered chondrocytes observed in osteoarthritis^47,48^. Here, cell clusters appear to manifest from supraphysiologic-TGF-β-induced hyperplasia, whereby proliferating chondrocytes are physically constrained in the agarose scaffold. Supraphysiologic TGF-β further reduces COL-II in the interterritorial matrix (Fig.6C), which may result from steric hindrance of collagen from sGAG-rich interterritorial regions^49^, or alterations to collagen structure and fibril architecture, leading to localization in the territorial regions^50,51^. This aberrant tissue organization may compromise mechanobiological cell signaling pathways needed for tissue homeostasis in response to physiologic loading. In contrast, LTGF-β scaffolds generate neocartilage that better resembles the morphology of native hyaline cartilage with more isolated chondrocytes and denser interterritorial collagen distribution. Based on the importance of cell-matrix interactions on cartilage homeostasis^52,53^, this improved tissue organization will likely yield important benefits to the maintenance of cartilage in its native chemical environment.

The long-term health of cartilage further requires cells to adopt a stable chondrogenic cell phenotype to provide the continued replenishment of hyaline cartilage matrix constituents (e.g., aggrecan and COL-II). Clinical cartilage regeneration platforms are commonly limited by a tendency of cells to produce fibrocartilage, as marked by enhanced COL-I expression^6,54,55^, or undergo hypertrophy, as marked by enhanced COL-X expression^56,57^. Here, this challenge is prominently illustrated as constructs cultivated in the absence of exogenous TGF-β exhibit enhanced COL-I expression (Fig.5F), while constructs cultivated with supraphysiologic TGF-β exhibit enhanced COL-X expression (Fig.5D), together suggesting that a stimulatory balance may be needed to maintain a chondrogenic phenotype. Physiologic TGF-β dosing via LTGF-β scaffolds provides this balance by maintaining both low COL-I (91-96% reduced relative to TGF-β-free) and COL-X (86-94% reduced relative to MS-10) expression, without detriment to the expression of COL-II, aggrecan, and SOX9 (Fig.5A-C). These data suggest that cells in neocartilage derived from LTGF-β scaffolds may be able to achieve improved long-term maintenance of the hyaline cartilage ECM. Interestingly, in prior work, supraphysiologic TGF-β was shown to increase COL-I protein deposition in neocartilage^15,58^, suggesting that transcription, translation, and post-translational regulation of collagen subtypes may be differentially regulated by TGF-β in chondrocytes^59,60^.

For small-size constructs, physiologic TGF-β doses, whether they are delivered via LTGF-β-conjugated scaffolds or alternatively through media supplementation of physiologic active TGF-β, improve tissue composition and cell phenotype. However, for constructs that approach the size required for clinical cartilage repair, TGF-β delivery via media supplementation proves inadequate due to the limited penetration of active TGF-β into the construct. As shown previously, these transport limitations result from high rates of cell-mediated internalization of the active TGF-β ligand, leading to significant active TGF-β gradients through the construct depth^19^. Consequentially, construct sGAG deposition and ECM stiffening are predominantly localized to the construct periphery—for physiologic dose supplementation, the deep tissue regions are ~6-fold softer than the media-exposed tissue periphery (Fig.7K-N). These results highlight the significant limitation of using media supplemented TGF-β to generate engineered cartilage for clinical use. In contrast, LTGF-β scaffolds overcome this limitation by delivering TGF-β via localized activation of LTGF-β within the construct. As a result, a large fraction of chondrocytes receives physiologic TGF-β doses, leading to a more uniform tissue development. Interestingly, residual mechanical property heterogeneities remain in LTGF-β constructs, suggesting that transport limitations of other nutrients and biomolecules may contribute. In future work, LTGF-β scaffolds will be incorporated with other biomolecule transport enhancing strategies^61^ to yield further improvements to neocartilage regeneration.

An additional consideration for TGF-β-incorporated scaffolds is the potential for TGF-β desorption from the scaffold post-implantation, leading to off-target effects on surrounding synovial joint tissues. Elevated levels of active TGF-β in synovial fluid are pathogenic, leading to fibrosis of the synovium and the emergence osteophytes^16,17^. Further, for scaffolds used in clinical bone regeneration, off-target effects from growth factor desorption have been associated with a myriad of complications and pathologies^62,63^. It remains unclear whether TGF-β releasing scaffolds in prior studies^9,12–14^ (e.g., heparin, microspheres, liposomes) release pathogenic levels of TGF-β into synovial fluid. However, given the highly supraphysiologic levels of active TGF-β loaded into these scaffolds, caution should be taken when evaluating the safety of TGF-β releasing scaffold products. Given these considerations, LTGF-β scaffolds may provide an additional benefit of mitigating off-target TGF-β effects on synovial joint tissues post construct implantation. While LTGF-β undergoes a degree of release from the construct over time, the prerequisite of activation of the latent complex to elicit cell signaling may provide a barrier that protects against off-target effects. Ultimately, LTGF-β may be cleared from the synovial joint prior to activation, thus mitigating off-target pathology.

The improved regenerative outcomes observed for LTGF-β scaffolds align with evolutionary mechanisms that exist to regulate TGF-β activity in native tissue^22^. Breakdown of these regulatory mechanisms yields a detrimental pathology, highlighting the importance of TGF-β regulation for native tissue development and maintenance^64,65^. A limitation of this study is the use of immature bovine chondrocytes that may activate LTGF-β at elevated rates or respond to subsequent physiologic dosing profiles differently than cell populations currently utilized in clinical practice. Future investigations will evaluate the efficacy of LTGF-β scaffolds in improving cartilage regeneration for human cell populations currently in clinical use or preclinical development, including chondrocytes, mesenchymal stem cells, induced pluripotent stem cells, and other chondrogenic progenitors. In summary, motivated by bio-inspiration, LTGF-β scaffolds provide an opportunity to overcome conventional TGF-β delivery limitations, giving rise to neocartilage with improved composition, structure, material properties, and cell phenotype.

## 5. Conclusions

In this study, we establish a bio-inspired LTGF-β conjugated scaffold to improve neocartilage development. The LTGF-β scaffold addresses the TGF-β delivery paradox in conventional media supplementation systems, where physiologic doses exhibit limited penetration and supraphysiologic doses induce aberrant tissue features. Scaffold-conjugated LTGF-β is uniformly distributed in tissue constructs and activated by embedded cells, leading to a local and moderate delivery of TGF-β. As a result, LTGF-β constructs achieved native-matched mechanical properties, significant enhancements in ECM compositions, and more hyaline-cartilage tissue features including dominant chondrogenic marker gene expressions, isolated cell morphology formation, and native-like ECM distributions. Further, scaffold-conjugated LTGF-β greatly reduces mechanical and biochemical heterogeneities in large-scale constructs approaching clinical use, which can benefit long-term maintenance of neocartilage post-transplantation. Together, the novel TGF-β delivery strategy via LTGF-β conjugation described in this work improves neocartilage development and is advantageous over current TGF-β supplemented media protocols.

## Author contributions

TW and MBA designed the study. MBA supervised the study and procured funding. TW, CCM, BH, EB, and AM performed experiments. TW, MDL, JEM, MWG, and MBA contributed to scientific discussions and data interpretation. All authors contributed to the preparation of the manuscript.

