## Supplementary Materials for "A Bio-inspired Latent TGF-β Conjugated Scaffold Improves Neocartilage Development"

**1. Effect of different TGF-β isoforms on construct growth.**

Methodology: Bovine chondrocyte-seeded agarose tissue constructs were cultured in chondrogenic media. Medium was supplemented with 10 ng/mL active TGF-β1 or 10ng/mL active TGF-β3 for the initial 2 weeks of culture. The compressive Young's modulus of constructs was measured after 38 days of culture. An additional group of constructs was cultured in the absence of TGF-β entirely (TGF-β free) as control.

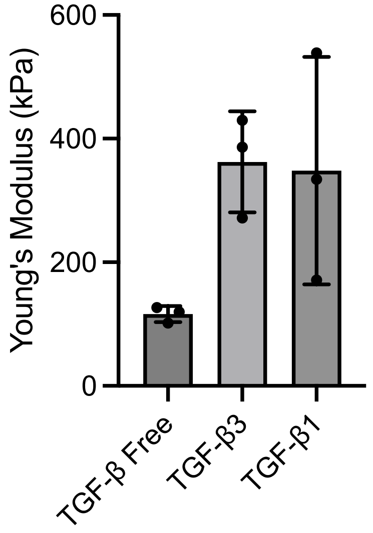


**Fig. S1** Compressive Young's modulus of cartilage constructs after 38 days of culture. Constructs exhibited similar enhancements in mechanical properties whether exposed to the TGF-β1 or TGF-β3 isoform in its active form. n=3. Error bars represent mean ± s.d.

**2. Effect of agarose methacrylate functionalization on construct growth.**

Methodology: Bovine chondrocytes were seeded in unmodified agarose (Agr) or methacrylate modified agarose (MeAgr) to fabricate tissue constructs. Constructs were cultured in chondrogenic media with or without 10 ng/mL active TGF-β3 for the initial 2 weeks of culture. Construct mechanical properties were measured at day 56.

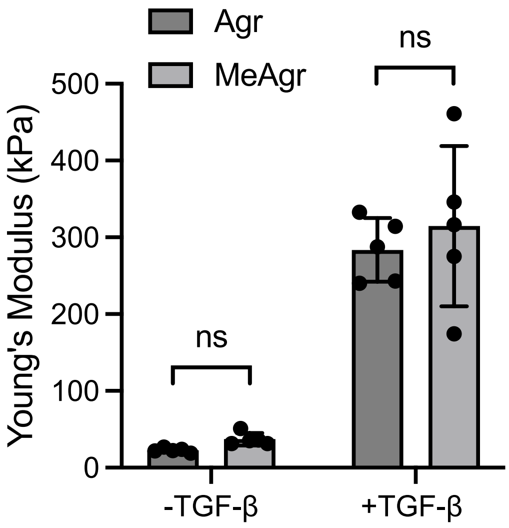


**Fig. S2** Day 56 compressive Young's modulus of cartilage tissue constructs fabricated with Agr or MeAgr ±TGF-β media supplementation. n=5. No significant differences in properties are observed for constructs fabricated from methacrylate functionalized agarose ($p>0.6$). Error bars represent mean ± s.d. ns: no significance.

**3. Effect of LTGF-β conjugation on cell viability.**

Methodology: Tissue constructs were fabricated from all TGF-β dosing groups: TGF-β-free, physiologic active TGF-β media supplementation (MS-0.3), supraphysiologic active TGF-β media supplementation (MS-10), and LTGF-β conjugation levels (Conj-Low, Conj-Med, Conj-High). After 42 days of culture, constructs were diametrically halved and stained with Live/Dead kit (Invitrogen). Images were acquired using confocal microscopy (Olympus FV3000, 10×).

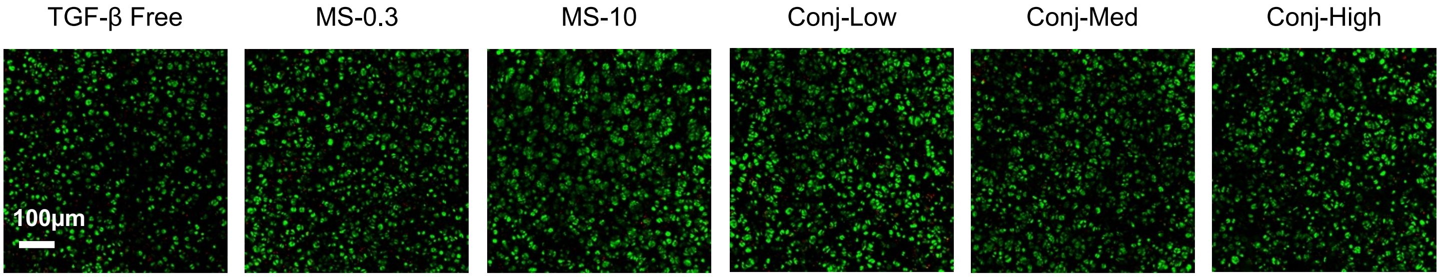


**Fig. S3** Confocal images for cell viability of tissue constructs with LTGF-β conjugation, active TGF-β media supplementation, or without TGF-β exposure. Green represents live cells and red represents dead cells. n=3.

**4. Covalent scaffold conjugation is required for LTGF-β-mediated enhancement of construct growth.**

Methodology: LTGF-β1 was loaded into methacrylate functionalized agarose (MeAgr) or nonfunctionalized agarose (Agr) at a dose of 3 μg/mL prior to bovine chondrocyte encapsulation. Tissue constructs were cultured for 42 days in the absence of media supplemented active TGF-β and analyzed for mechanical properties, sGAG content, and collagen content. An additional group of constructs was fabricated in Agr without initial LTGF-β loading and cultured in the absence of media supplemented active TGF-β (TGF-β free) as a control.

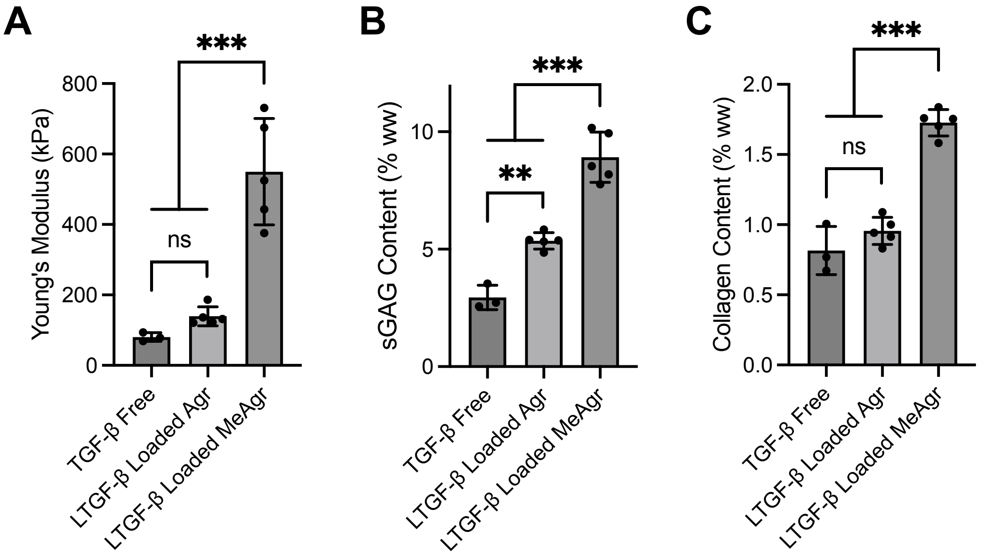


**Fig. S4** (**A**) Compressive Young's modulus, (**B**) sGAG content, and (**C**) collagen content (**C**) of cartilage tissue constructs cultivated for 42 days with LTGF-β initially loaded into a methacrylate functionalized or unfunctionalized agarose scaffold. Mechanical properties and biochemical contents are significantly elevated for constructs fabricated from methacrylate functionalized agarose, indicating that chemical conjugation of LTGF-β to the scaffold is required for LTGF-β mediated enhancements of tissue growth to occur. n=5. Error bars represent mean ± s.d. **p<0.005, ***p<0.001, ns: no significance.
